# High-Throughput Evolution Unravels Landscapes of High-Level Antibiotic Resistance Induced by Low-Level Antibiotic Exposure

**DOI:** 10.1101/2023.11.30.569484

**Authors:** Hanqing Wang, Hui Lu, Chao Jiang, Lizhong Zhu, Huijie Lu

## Abstract

Potential pathogens consistently exposed to low-level environmental antibiotics could derive high-level clinically relevant resistance detrimental to the human health. However, the underlying evolutionary landscapes remain poorly understood. We conducted a high-throughput experimental evolution study by exposing an environmentally isolated pathogenic *Escherichia coli* strain to 96 typical antibiotics at 10 μg l^−1^ for 200 generations. Antibiotic resistance phenotypic (IC_90_ against 8 clinically used antibiotics) and genetic changes of the evolved populations were systematically investigated, revealing a universal increase in antibiotic resistance (up to 349-fold), and mutations in 2,432 genes. Transposon sequencing was further employed to verify genes potentially associated with resistance. A core set of mutant genes conferring high-level resistance was analyzed to elucidate their resistance mechanisms by analyzing the functions of interacted genes within the gene co-fitness network and performing gene knockout validations. We developed machine-learning models to predict antibiotic resistance phenotypes from antibiotic structures and genomic mutations, enabling the resistance predictions for another 569 antibiotics. Importantly, 14.6% of the 481 key mutations were observed in clinical and environmental *E. coli* isolates retrieved from the NCBI database, and several were over-represented in >500 clinical isolates. Deciphering the evolutionary landscapes underlying resistance exposed to low-level environmental antibiotics is crucial for evaluating the emergence and risks of environment-originated clinical antibiotic resistance.

## Introduction

The emergence of antimicrobial resistance (AMR) in the environment is a growing concern in the One Health paradigm^1–5^. Antibiotics are increasingly released into the environment via wastewater discharge and disposal of solid waste, where 40-90% are excreted as parent compounds^6^. Environmental bacteria exposed to these antibiotics at low-level, sub-minimal inhibitory concentrations (sub-MIC) can evolve antibiotic resistance by accumulating diverse low-frequency mutants, which are different from those evolved at clinical concentrations^7–11^. The environmental research community essentially needs to elucidate the contribution of mutation-driven resistance evolution in the environment by crafting analytical measures to pinpoint the responsible selective forces (i.e., antibiotics) driving AMR development^12^. Therefore, exploring the evolutionary landscape under low-level antibiotic exposures is increasingly important at the interface between environmental and clinical microbiology pertaining to antibiotic resistance^4,13^.

Traditional low-throughput experimental evolution has provided an incomplete view of the AMR evolutionary landscape, as the number of stress conditions, mutants, and resistance-conferring genes examined were limited^14,15^. High-throughput experimental evolution enables the rapid generation and systematic investigation of population-level antibiotic resistance under a variety of stress conditions, allowing better characterization of the structure- or dose-response relationships among stressors and induced resistance^16,17^. In addition, antibiotic resistance genes (ARGs) were traditionally identified by sequencing and verified through phenotypic assays/knockout experiments, bearing the risks of overlooking critical insights from genetic interactions. Transposon sequencing (Tn-Seq) measures genome-wide gene fitness across large-scale transposon mutagenesis libraries. By combining Tn-Seq with deep resequencing, researchers can map the accumulated mutations onto complex and coordinated interaction networks. The prevalence and clinical significance of identified mutations in broad environments can be examined by comparative genomics^18^. The above novel approaches could help address critical knowledge gaps in the field of environmental AMR, for example: (1) What is the association between the antibiotic resistance of evolved populations and the chemical structures of antibiotics under low-level exposures? (2) What cellular functions are affected by mutations potentially via new antibiotic resistance mechanisms? (3) Are mutations that arise under low-level antibiotic exposures observed in clinically isolated resistant strains?

This study utilized a environmental opportunistic pathogen *E. coli* strain to perform high-throughput experimental evolution under low-level exposures to 96 antibiotic stressors (**Supplementary Fig. 1**). The antibiotic resistance profiles of evolved populations and their core mutations were characterized. To explore the resistance mechanisms conferred by these core mutant genes, a genome-wide co-fitness interaction network was constructed based on Tn-seq. Chemical fingerprint-based modeling was built to predict antibiotic resistance profiles induced by another 569 antibiotics sharing similar structural features with the 96 stressors. The occurrence of core mutations in broader environmental and clinical isolates was examined by comparative genomics. These findings underscore the importance of AMR developed under low-level antibiotic exposures in environmental settings and its potential spread to clinical contexts.

## Results

### Antibiotic resistance profiles of evolved *E. coli* QSHQ1 populations

*E. coli* strain QSHQ1 isolated from the environment evolved under exposure to 96 antibiotic stresses at an environmentally relevant concentration of 10 μg l^−1^ for 20 days, which corresponded to approximately 200 generations (**Supplementary Data 1**). These antibiotics can be classified into ten classes and mainly act through four mechanisms, including inhibition of DNA/RNA (DSI), protein (PSI), cell wall (CWSI), and membrane (MSI) synthesis (**Fig. 1a**). All populations survived and displayed varying degrees of growth inhibition compared to control populations without antibiotics (**Supplementary Fig. 2**). Evolved populations on day 20 were tested for antibiotic resistance (indicated by IC_90_) to eight clinically relevant antibiotics, including azithromycin (AZI), ciprofloxacin (CIP), tetracycline (TET), sulfamethoxazole with trimethoprim (SMZ+TMP), tobramycin (TOB), ceftazidime (CAZ), meropenem (MER), and polymyxin B (PMB). These IC_90_ values were compared with CLSI’s breakpoint values for susceptible and resistant bacteria (**Fig. 1b** and **Supplementary Data 2**).

**Fig. 1:**
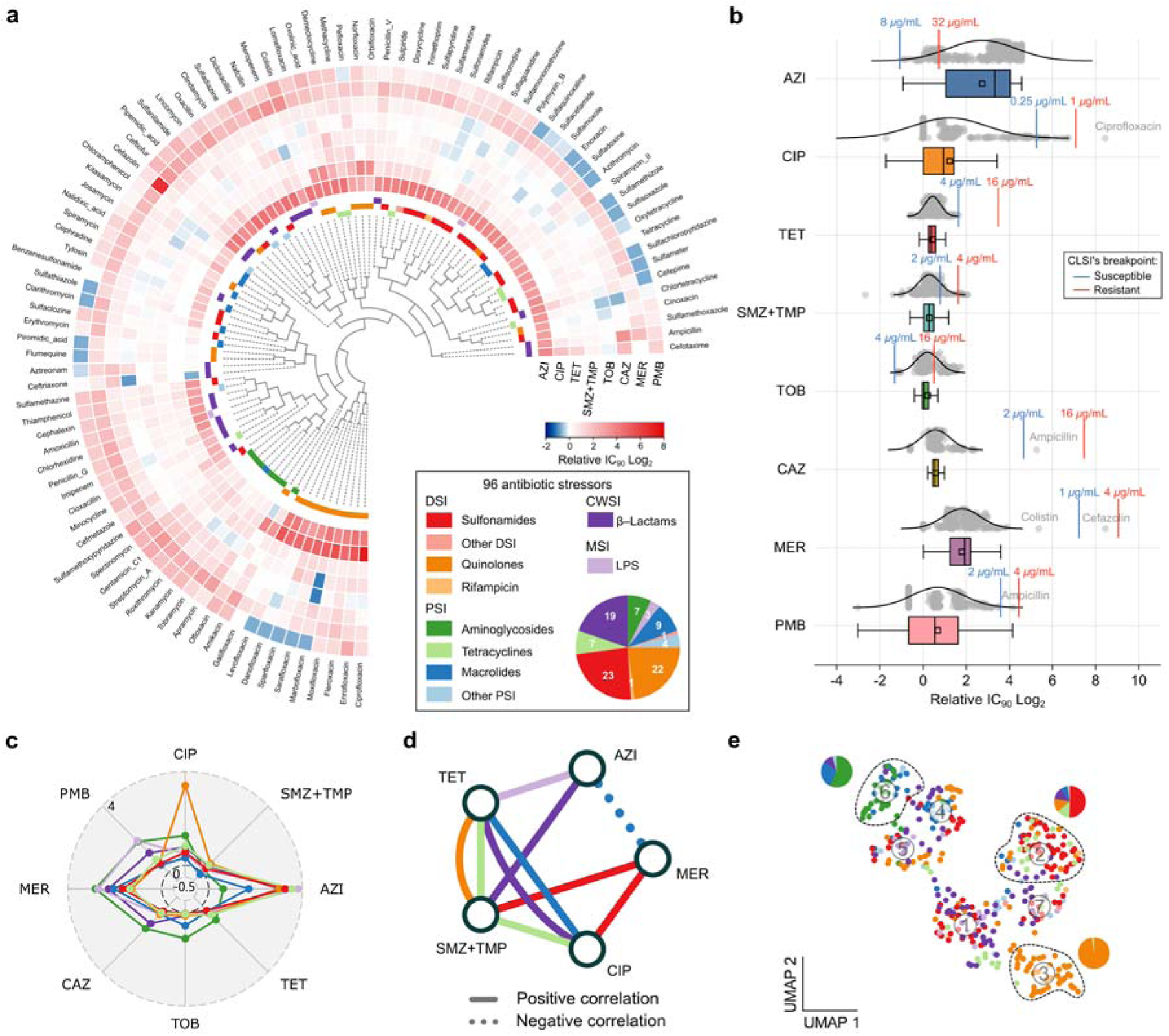
Antibiotic resistance profiles of evolved populations. **a,** Antibiotic resistance profiles of the evolved populations. Antibiotic resistance changes were indicated by log_2_-transformed fold changes in IC_90_ values (relative to the control populations without antibiotic stress). The tree was constructed based on Euclidean distances. Antibiotics stressors: 96; Stressor classes: 10 (different colors of the inner circle); Stressor groups: 4 (DSI, CWSI, PSI, and MSI); Antibiotic resistance assay: 8 (heatmap circles); Biological replicates: 4. **b,** Distribution of antibiotic resistance changes in evolved populations. Grey dots: log_2_-transformed fold changes in IC_90_ values of all independently evolved populations; Curves: normal distribution approximation; Colored sticks: susceptible (blue) and resistant (red) breakpoints for the tested antibiotic resistance according to CLSI guidelines^21^. Box plot: center line, median; small square, mean; left and right sides of the box, upper and lower quartiles; whiskers, 1.5× interquartile range. **c,** Averaged resistance changes (log_2_-transformed fold changes in IC_90_ values) of evolved populations stressed by the same class of antibiotics. **d,** Positive and negative correlations between different antibiotic classes according to the antibiotic resistance profiles (Pearson’s correlation coefficient |*r|* > 0.6, *P* < 0.05). **e,** Uniform manifold approximation and projection (UMAP) embedding of evolved populations based on their resistance to eight antibiotics. **c**, **d**, and **e** are assigned the same color code as **a** for antibiotic classes.

The wild-type (WT) strain was initially susceptible or intermediate to the eight clinically relevant antibiotics. All 96 evolved populations conferred resistance (increased IC_90_) to at least one of the eight antibiotics (Multiple unpaired *t*-tests, FDR < 0.1, **Fig. 1a**), where resistance to AZI (78.9%) and TOB (18.8%) were the most prevalent. A similar evolution of AZI-resistant strains was recently reported for *Comamonas testosteroni*^10^. Only two evolved populations stressed by marbofloxacin and ceftriaxone displayed significantly decreased IC_90_ values to at least one antibiotic. Specifically, a ciprofloxacin-stressed population displayed the most significant increase in CIP resistance (IC_90_ increased by 345-fold), and a cefazolin-stressed population displayed a 349-fold increase in MER resistance (**Fig. 1b**). Previous studies^19,20^ reported 256-fold and >300-fold increase in the MIC for ciprofloxacin of *E. coli* and *S. aureus* cells after < 20 passages, respectively. In fact, the WT strain was susceptible to CIP and MER. Exposure to quinolones was the primary driver of CIP resistance evolution (average IC_90_ fold-change: 24.6 ± 14.9), while exposure to aminoglycosides highly contributed to the evolution of TOB resistance (average IC_90_ fold-change: 2.19 ± 0.26) (**Fig. 1c**, **Supplementary Figs. 3** and **4**).

Populations evolved under each class of antibiotics displayed variations in resistance to the eight antibiotics tested, where 5/8 displayed either significantly positive or negative correlations induced by a specific class of antibiotics (**Fig. 1d** and **Supplementary Fig. 5**). Overall, positive correlations (Pearson’s *r* > 0.6, *P* < 0.05) were much more abundant than negative ones, where the latter was only observed for macrolides-stressed populations and between their AZI and MER resistance (*r* = −0.85, *P* < 0.05). The 96 antibiotic stressors were further clustered based on the antibiotic resistance profiles they induced (**Fig. 1e** and **Supplementary Fig. 6**). Most aminoglycosides and macrolides (both acting via PSI) were clustered (VI), as well as sulfonamides and tetracyclines (acting via DSI and PSI, respectively, cluster II). Quinolones formed their distinct cluster (cluster III), separated from other antibiotic classes.

### Phenotypic convergence and divergence of antibiotic resistance evolution

Pairwise antibiotic stressors induced convergent (Spearman’s correlation coefficient ρ > 0.2, 76%) and divergent (ρ < −0.2, 4.6%) antibiotic resistance phenotypes among the 96 antibiotics (**Figs. 2a** and **2b**). Convergent phenotypes (a phenomenon whereby developing similar phenotypes) were predominant and occurred more frequently within the same class of stressors (85.3% vs. 73.9%, **Fig. 2c**). Divergent phenotypes (orthogonal), otherwise, were more found across different antibiotic classes than within the same class (5.1% vs. 2.3%). Overall, the frequency of convergence was much higher than the previously reported value (60%) for *P. aeruginosa* strains that evolved under high-level antibiotic exposures^22^. We hypothesize that the survival of more diverse resistant subpopulations under low-level antibiotic exposures likely contributes to widespread elevated population antibiotic resistance, as these subpopulations may possess similar off-target mutations^23^. However, the consistency of the mutation similarity and convergent phenotypes await confirmation via deep sequencing. Furthermore, stressors within aminoglycosides and lipopeptides (LPS) induced the most convergent resistance phenotypes, while quinolones induced the most divergent resistance patterns compared to other antibiotic classes (**Fig. 2d**). Convergent resistance phenotypes were widespread across classes. Divergent phenotypes were more frequently observed between aminoglycosides-quinolones and aminoglycosides-tetracyclines, e.g., kanamycin and sparfloxacin (ρ = −0.81; *P* = 0.0149) than other pairwise combinations (**Fig. 2e** and **Supplementary Fig. 7**). Our analyses of these stressor-stressor pairs corroborated the previous findings of resistance evolution leading to phenotypic convergence and identified new population-level convergent resistance evolution in sub-MIC antibiotic selections^22^.

**Fig. 2:**
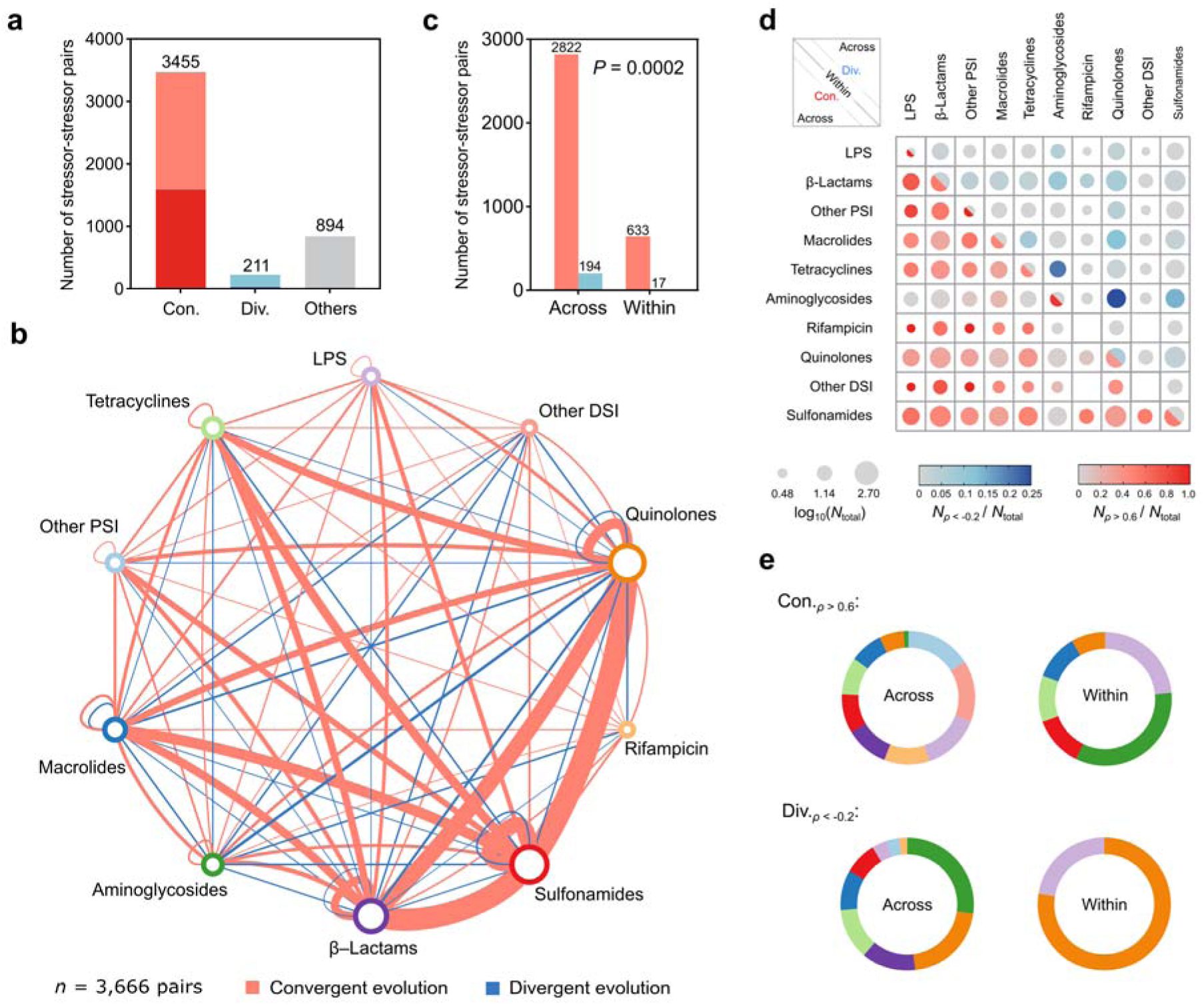
Convergent and divergent evolution relationships based on antibiotic resistance profiles induced by low-level antibiotic exposures. **a,** Total number of convergent (con.) and divergent (div.) evolution relationships. Strong convergent or divergent evolution is presented in dark red or blue, respectively (Spearman’s rank correlation |ρ| > 0.6); light red or blue suggests weak convergent or divergent evolution (0.2< |ρ| <0.6). **b,** Convergent and divergent evolution relationships between stressors belonging to different (across) or same (within) antibiotic classes. Node size reflects the number of stressors within each antibiotic class. Edge thickness reflects the number of convergent or divergent evolution pairs (|ρ| > 0.2). **c,** Convergent and divergent resistance evolution occurred between stressors across and within classes. Convergent: red; divergent: blue. *P* value: chi-square test significance. **d,** Frequency of convergent evolution (strong, ρ > 0.6, red) and divergent evolution (ρ < −0.2, blue) relationships among antibiotic classes. Dot size reflects the number of stressor pairs within (diagonal) or across (upper and lower triangles) any two antibiotic classes. **e,** Proportions of convergent and divergent evolution relationships ‘across’ and ‘within’ antibiotic classes. **e** is assigned the same color code as **b** for antibiotic classes.

### Mutations in evolved populations

For each stress condition, the four evolved cultures were pooled for deep resequencing (coverage: 1,642 ± 466, *n* = 97, including one control group without antibiotic exposure). A total of 56,744 mutation events were detected by filtering the mutations in the control group, averaging 591 mutations per condition (**Supplementary Fig. 8** and **Supplementary Data 3**), causing 4,750 amino acid changes (AACs) involved in 2,432 protein-coding genes (**Fig. 3a**). These mutations were classified into single nucleotide polymorphisms (non-synonymous SNPs, 3,179), insertions (379), and deletions (1,428). Here, we confirmed the consistency of convergent phenotypes with similar mutational backgrounds (**Fig. 3b**). Additionally, all antibiotic stressors selected more non-synonymous than synonymous substitutions (d*N*/d*S* > 1, positive selection), and there were negative correlations between d*N*/d*S* and the number of point mutations (**Fig. 3c**). Overall, DSIs induced more point mutations than other stressors at the same d*N*/d*S* level, consistent with their mechanisms of action.

**Fig. 3:**
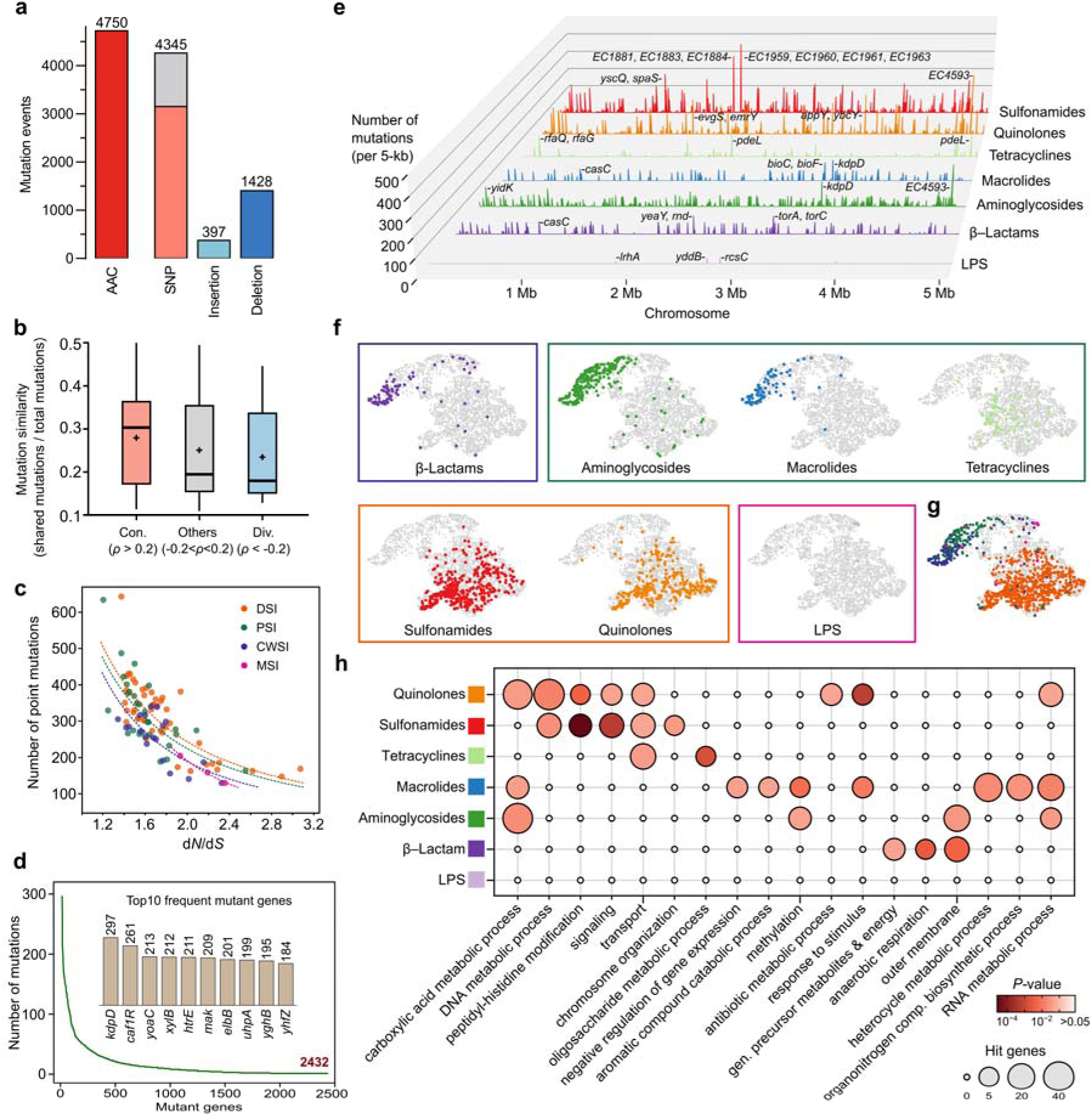
Mutations detected in evolved populations. **a,** Distribution of mutation events. SNPs are classified into non-synonymous (red) and synonymous (grey) ones. **b,** Mutation similarity in evolved populations exposed to stressors exhibiting convergent or divergent evolutionary relationships. Box plots: center line, median; ‘+’, mean; limits, first and third quartiles; upper and lower whiskers, the highest and lowest values within 1.5× the interquartile range from the hinge. Mutation similarity levels are significantly different for evolved populations exhibiting convergent, other, and divergent relationships (F = 44.59, *P* < 0.0001; one-way ANOVA, Tukey’s honestly significant difference). **c,** Relationships between non-synonymous to synonymous mutation ratio (d*N*/d*S*) and the number of point mutations. Dashed curves represent non-linear regressions performed for the four groups of stressors (*R*^2^ = 0.59-0.99, all *P* < 0.05). **d,** The distribution of mutation counts for a total of 2,432 mutant genes. The top ten mutant genes harboring the most mutations are displayed. **e,** Mutation hotspots in the chromosomes of evolved populations under different antibiotic stresses. The top three mutation hotspots are highlighted for each antibiotic class. **f,** UMAP clustering of highly variable AACs (*n* = 2,718) occurred under the stresses of seven antibiotic classes, which are further grouped based on the action mechanisms of antibiotics (four groups, rectangle frames). **g,** UMAP clustering of highly variable AACs among the four groups of stressors. **h,** GO biological processes associated with mutant genes for different classes of antibiotic stresses.

The mutation counts of the 2,432 genes exhibit a power-law distribution with a long tail, where the top ten mutant genes account for 6% of the total mutations (**Fig. 3d**). In order to find mutation hotspots along the chromosomal DNA, we performed mutation enrichment analysis in 5-kb sliding windows. The top three mutation hotspots were flagged for the seven classes of antibiotic stressors (**Fig. 3e**). Some sites with frequent mutations included, e.g., the acid stress response gene *evgS* and the multidrug efflux pump gene *emrY*. Interestingly, *kdpD* mutation frequency was relatively high in both macrolides and aminoglycosides-stressed populations. Recent studies report that KdpD promotes resistance to osmotic, oxidative and antimicrobial stresses^24^. *yeaY* with frequent mutation under β-lactam stressors encodes a putative outer membrane lipoprotein^25^. *rcsC* flagged under LPS stress is a known ARG that exhibited increasing expression in response to polymyxin B. Polymyxin E can activate the RcsC/YojN/RcsB two-component system, responding to alterations in the bacterial membrane^26^.

Unsupervised clustering UMAP embedding was used to test the hypothesis that antibiotics with the same mechanism of action would enrich mutations leading to similar AACs (**Fig. 3f**). The hypothesis was largely verified with one exception-tetracyclines (PSI), which enriched AACs similar to those enriched under sulfonamides and quinolones stresses (DSI) rather than aminoglycosides and macrolides (PSI). This aligned well with the observed antibiotic resistance phenotypes stressed by these antibiotics. Tetracyclines could likely induce DNA damage due to their strong binding affinity to DNA and their ability to generate free radicals in close proximity to the DNA backbone^27,28^. Among the 2,718 highly variable AACs, 991 were enriched by at least one of the four action mechanisms of stressors (Fisher’s exact test *P* < 0.05). DSI antibiotics contributed the most to the overall enrichment (722 hits), and the majority of overlapping hits were observed between PSI and CWSI antibiotics (**Fig. 3g**).

We next used Gene Ontology (GO) enrichment analysis to understand the co-occurrence of mutant gene-associated biological processes for the seven classes of antibiotics (**Fig. 3h**). ‘DNA metabolic process’ was represented among the top hits by sulfonamides and quinolones (DSI), and ‘outer membrane’ was the most targeted gene category by LPS stressors (MSI). Meanwhile, tetracyclines caused the most mutant genes in the ‘transporter’ category, including but not limited to those related to carbohydrate transport and metal ion transport. Methylation of the 16S rRNA is an emerging mechanism of high-level resistance to PSI antibiotics via ribosomal methyltransferases^29,30^. Mutant genes associated with this process were significantly more prevalent in the presence of macrolides and aminoglycosides compared with other antibiotics.

### Mutant resistance-relevant genes and co-fitness interaction network

Recognizing that not all mutant genes directly contribute to high-level resistance, we employed Tn-Seq to determine gene essentiality and identify potentially resistance-relevant genes (PRGs) across the whole genome. Successful construction of the Tn library using the WT strain enabled high-confidence essentiality and fitness quantification in the presence and absence of eight antibiotics used in MIC assays (**Supplementary Fig. 9**). Overall, the WT strain genome contained 468 essential, 3,910 non-essential, and 367 uncertain genes (**Fig. 4a**). Genes with absolute fitness change |Δ*W|* > 0.15 for at least one antibiotic were considered as PRGs (*n* = 2,830, **Fig. 4b** and **Supplementary Data 4**). Their proportions increased from 55% across the whole genome to approximately 73% among mutant genes (**Fig. 4a**). This highlighted that PRGs were indeed selected during experiment evolution under antibiotic stresses.

**Fig. 4:**
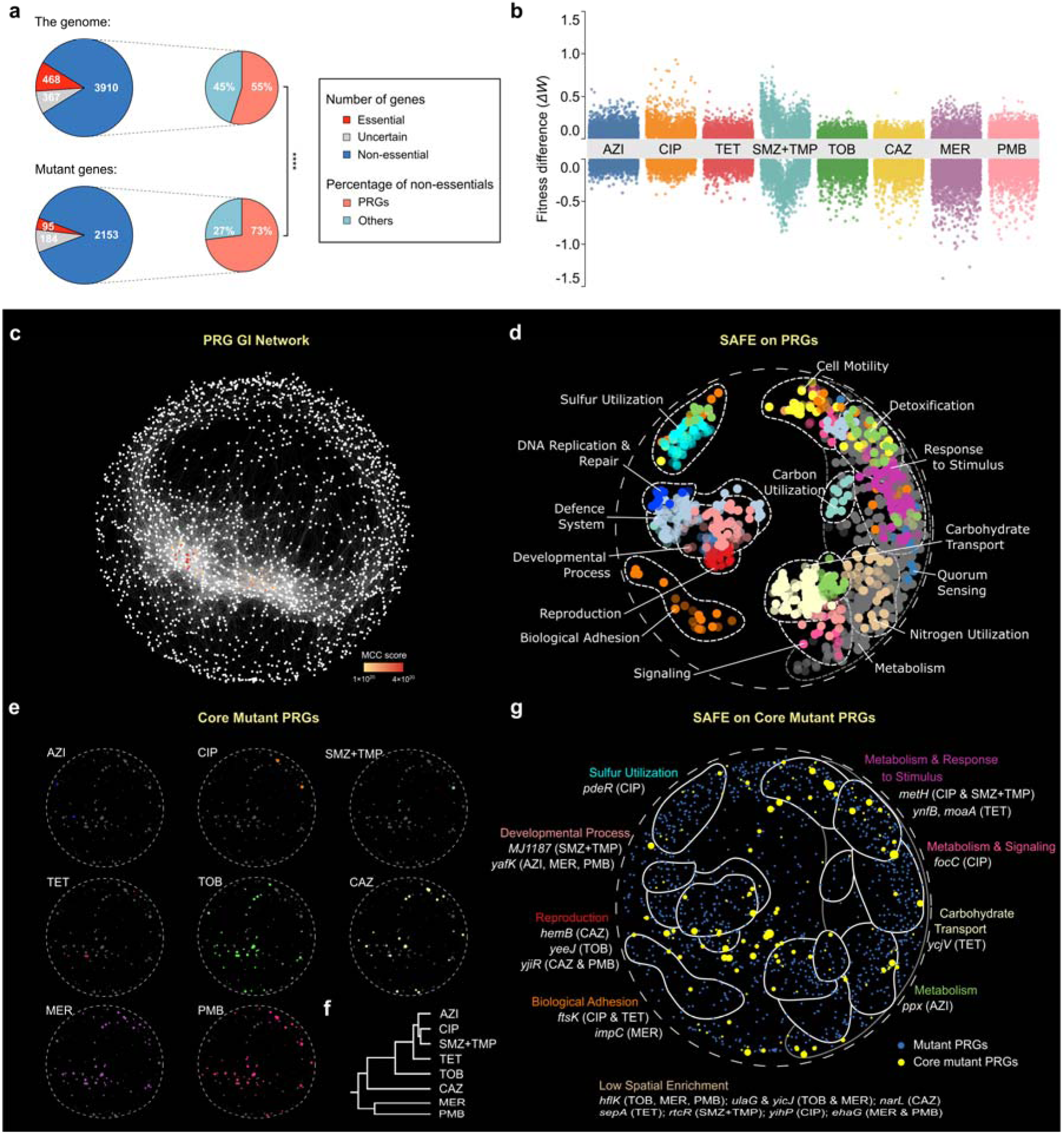
Gene co-fitness interaction network and spatial analysis of functional enrichment. **a,** Number of essential, non-essential, uncertain genes in the whole genome and mutant genes. ****: chi-square test *P* < 1*e*–4. **b,** Fitness differences (Δ*W*) of all Tn-Seq library genes for the 8 tested antibiotics. **c,** Gene interaction (GI) network of PRGs (*n* = 2,152) based on their co-fitness matrix. **d,** Enriched 15 bioprocesses by SAFE analysis on the PRG GI network in **c**. **e,** Core mutant PRGs conferring high-level resistance to the eight antibiotics (nodes, *n* = 131). Node size reflects the co-occurrence frequency of core mutant PRGs. **f,** Hierarchical clustering of the eight antibiotics based on their associated core mutant PRGs. **g,** Mutant PRGs (blue) and core mutant PRGs (yellow) are highlighted on a schematic representation of the GI network. The 22 labelled representative core mutant PRGs are associated with SAFE high-enrichment bioprocesses (within the circle, *P* < 0.05, corrected for multiple testing) or low-enrichment bioprocesses (outside the circle).

To map the co-fitness interactions of pairwise PRGs, a 2,830 × 2,830 correlation matrix was calculated based on their conditional fitness profiles. A total of 22,811 co-fitness interactions were identified for 2,152 PRGs (Pearson’s |*r|* > 0.75, corrected *P* < 0.00002), including 15,403 positive and 7,408 negative interactions that were used for constructing the high-confident co-fitness interaction network (**Fig. 4c**). Highly connected hub PRGs (Maximal Clique Centrality, MCC score > 1*e*20) were identified, forming two central regions in the interaction network. Spatial analysis of functional enrichment (SAFE) further identified 15 network regions spatially enriched for similar bioprocesses, encompassing 1,093 PRGs in total (high-enrichment PRGs, **Fig. 4d** and **Supplementary Data 5**). In addition to some well-studied resistance-associated bioprocesses such as DNA replication and repair, response to stimulus, adhesion, and metabolism, the SAFE analysis also uncovered the enrichment of bioprocesses previously unreported (e.g., carbon/nitrogen/sulfur utilization) or less known (e.g., metabolism or cell motility^31,32^) to be associated with antibiotic resistance. How these cellular bioprocesses influence resistance evolution under low antibiotic stresses warrants further investigation.

Mutant PRGs (*n* = 1,572) that potentially contribute to the high-level antibiotic resistance of evolved populations, i.e., core mutant PRGs (*n* = 131, *P* < 0.01, Fisher’s exact test), were identified by comparing those enriched in populations exhibiting the top 10 versus bottom 40 resistance increase (**Supplementary Fig. 10**). The majority of core mutant PRGs were linked to high-level resistance against TOB, CAZ, MER, and PMB (**Fig. 4e, 4f,** and **Supplementary Data 6**). A representative subset comprising 22 out of the 131 core mutant PRGs was selected for more detailed investigation on their mechanisms of resistance contribution, including 9 hub genes identified in **Fig. 4c**. Some of these representative core mutant PRGs participate in bioprocesses that were spatially enriched by SAFE analysis (high enrichment, **Fig. 4g**), such as response to stimulus (e.g., *ynfB*^33^), signaling (*focC*^34^), and carbohydrate transport (*ycjV*^35^). While other core mutant PRGs contribute to bioprocesses that were not spatially enriched (low-enrichment), such as transcriptional reprogramming genes (e.g., *rtcR* and *narL*), they may still play roles in the development of AMR. This is likely because compared with high-enrichment PRGs, the low-enrichment ones exhibit higher degrees of co-fitness interactions (**Supplementary Fig. 11**), indicating their potential to interact with broader PRGs and thereby facilitate AMR development.

### Resistance mechanisms of representative core mutant PRGs

Among the 22 representative core mutant PRGs, 16 have reported links to specific resistance mechanisms, including biofilm formation (e.g., *focC* and *pdeR*), efflux pump (e.g., *ycjV*), drug inactivation (e.g., *MJ1187*), target bypass (e.g., *yafK),* etc. We are therefore interested in uncovering the contribution mechanisms of the remaining 6 representative core mutant PRGs with unreported links to resistance (i.e., *metH, ynfB*, *ulaG, ftsK*, *rtcR*, and *hflK*) by constructing their gene interaction modules (module I-VI, respectively, **Fig. 5a**). Within their modules, these PRGs interacted with genes encoding biofilm formation, metabolism, efflux pump, target alteration, membrane permeability, and DNA replication & repair-related proteins. On the one hand, these connections indicated that the 6 PRGs likely contribute to AMR development through similar mechanisms. On the other hand, analyzing the fitness differences (Δ*W*) of PRG-connected genes in response to antibiotic exposure further refined the proposed resistance mechanisms (**Fig. 5b**). For example, the fimbrial adhesin protein-coding gene *faeJ* and biofilm formation associated gene *vasD*, which interacted with *rtcR* (module V), exhibited significant fitness differences (|Δ*W| =* 0.30 ± 0.05 under SMZ+TMP stress). Similarly, the interactions of *hflK* (module VI) with the fimbrial adhesin protein-coding genes *yehB*, *afaA*, and biofilm formation associated gene *suhB* (|Δ*W| =* 0.35 ± 0.19 in response to TOB, MER, and PMB stresses) suggested that *hflK* may also contribute to AMR development through affecting biofilm formation.

**Fig. 5:**
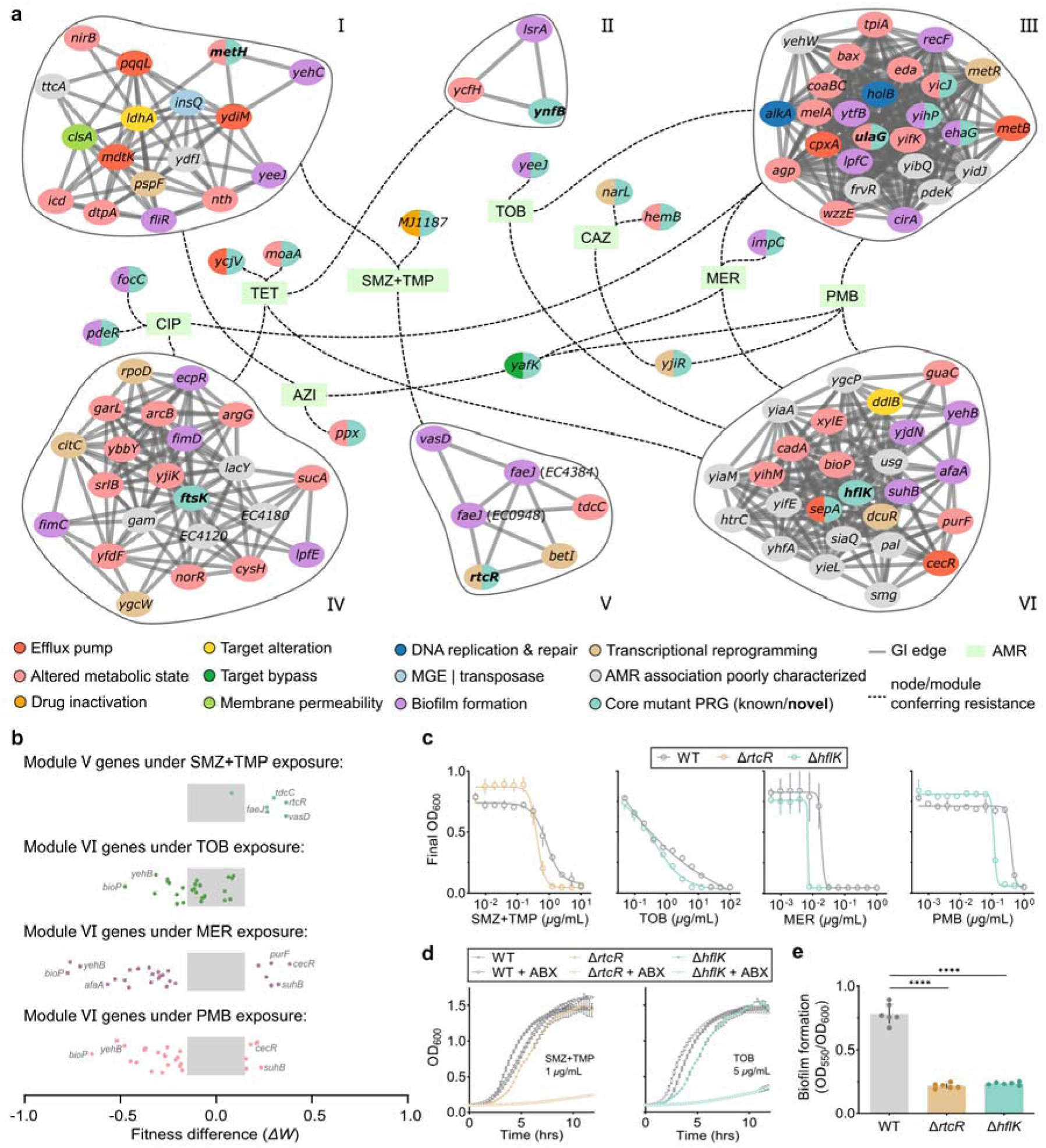
Gene co-fitness interaction modules of selected core mutant PRGs and verification of their proposed resistance mechanism. **a,** Core mutant PRGs displayed within modules extracted from the gene co-fitness interaction network (in bold, with unknown resistance mechanism) or outside modules (with known resistance mechanisms). **b,** Fitness differences of genes in modules V and VI under specific antibiotic exposures. Grey regions represent abs(Δ*W*) < 0.15. **c,** Dose–response curves for the three strains exposed to antibiotics at varied concentrations. Error bars represent standard deviations from biological triplicate. **d,** Decreased resistance of Δ*rtcR* and Δ*hflK* strains to SMZ+TMP and TOB, respectively, compared with the WT strain. **e,** Biofilm formation potentials of the WT, Δ*rtcR*, and Δ*hflK* strains determined by the crystal violet assay. *N* = 6 for all strains. ****: *P* < 0.0001.

The proposed resistance mechanisms of *rtcR* and *hflK* were further validated via gene knockout experiments. The two knockout strains exhibited increased susceptibility to their respective antibiotics (**Fig. 5c**), prolonged lag phases in growth (*P* < 0.05, Student’s *t*-test; **Fig. 5d**), and decreased biofilm formation (approximately 72% lower than that of the WT strain, **Fig. 5e**). These results collectively suggest that *rtcR*^M^ and *hflK*^M^ may confer resistance by enhancing biofilm formation, thereby limiting antibiotics diffusion into the population. The resistance mechanisms of the remaining four representative core mutant PRGs (*metH, ynfB*, *ulaG,* and *ftsK)* could be proposed and validated using similar approaches in future studies.

### Resistance evolution prediction by machine-learning models

A long-standing challenge in AMR research has been to predict the outcome of resistance evolution based on the chemical structure and/or dose of stress compounds. The 96 antibiotics used in this study are adequately representative of the structure diversity of commonly applied antibiotics. We developed machine learning models that can correlate the chemical structures of antibiotics, represented by extended-connectivity fingerprints (ECFPs), with the antibiotic resistance profiles of evolved strains. Automated generations by genetic algorithms were employed for feature selection and model optimization. The ensembled regression models demonstrated varied modeling performance on the training datasets, with coefficient of determination (*R*^2^) values ranging from approximately 0.3 to 0.8 for the eight antibiotics (**Fig. 6a**). The modeling performance on CIP resistance (*R*^2^ = 0.79 ± 0.01) was better than those for other antibiotics (F = 31.48, *P* < 1*e*–4, two-way ANOVA). We then used CIP resistance prediction results as an example to understand the contributions of monomer structures to antibiotic resistance. The top five antibiotics leading to high-level CIP resistance changes (log_2_ fold-change from 5.16 to 7.32) were annotated for their atomic groups’ contributions to the CIP resistance (**Supplementary Fig. 12)**. The key contributors to CIP resistance changes were the fluorine atom introduced at the C-6 position and the cyclopropyl group introduced at the N-1 position of the parent nucleus.

**Fig. 6:**
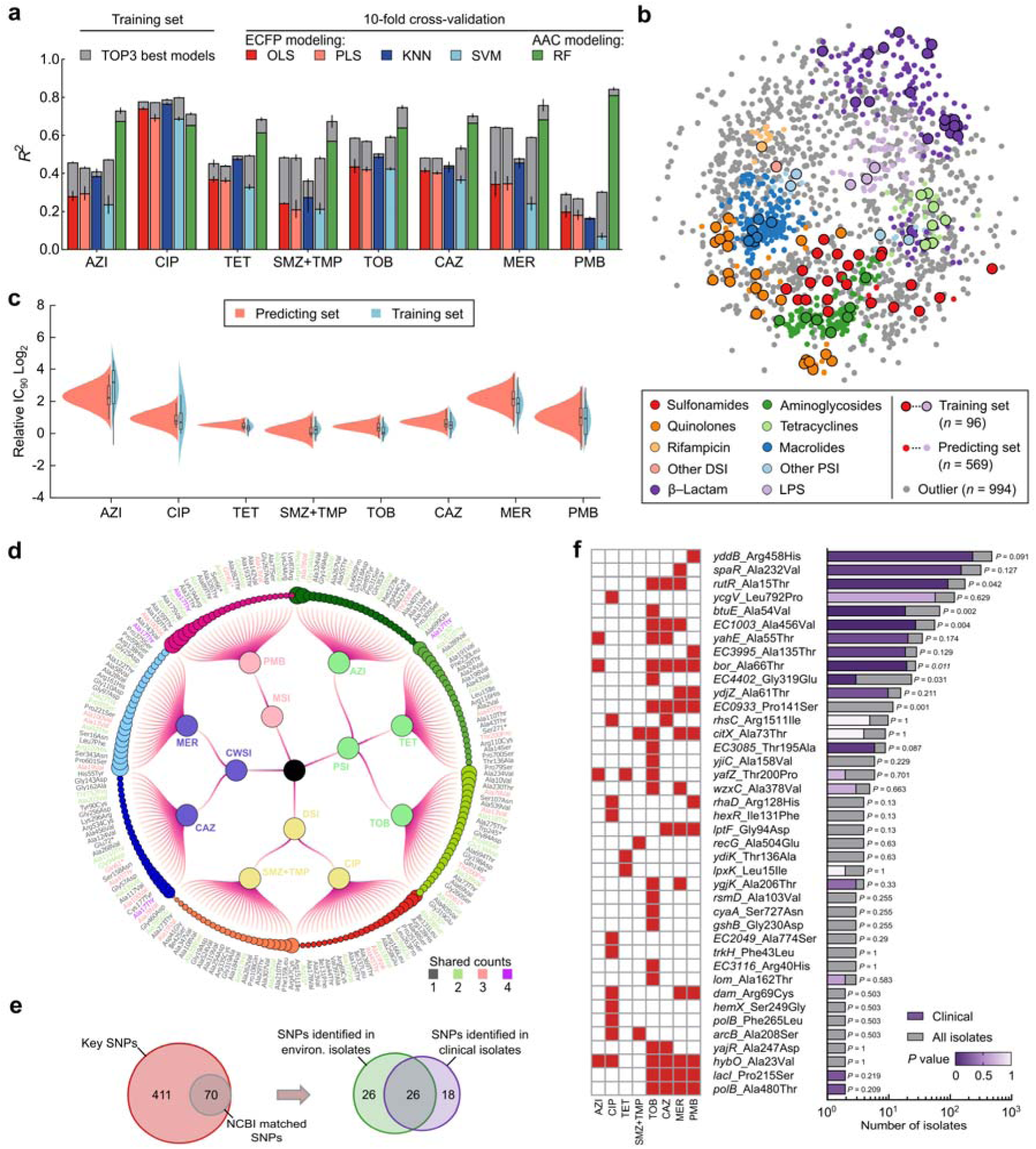
Performance of machine learning models and prevalence of key point mutations in clinical/environmental isolates. **a,** Performance of the machine learning models. The error bars represent the standard error of mean from the top three best-performing models. **b,** Tanimoto similarity network demonstrating the applicability domain (Tanimoto similarity ≥ 0.2) of ECFP-based models. **c,** Split violin plots showing the normalized distributions of predicted resistance levels for the training (*n* = 96) and predicting antibiotics (*n* = 569). **d,** Top 25 most important SNPs (features) contributing to the accuracy of the AAC-based RF model for each antibiotic. Dot size: importance of features; Shared counts: occurrence of particular SNPs in the circle. **e,** Occurrence of key SNPs (feature-importance SNPs in **d** and SNPs in core mutant PRGs conferring high-level resistance, *n* = 481 in total) in NCBI-retrieved clinical and environmental isolates. **f,** Left: key SNPs associated with resistance to the eight antibiotics. Right: number of clinical and environmental isolates containing the top 40 key SNPs. Total isolates: 1,263 retrieved from the NCBI Pathogen Detection database. Fisher’s exact test *P* values indicate the overrepresentation of SNPs in clinical isolates and are colored in varying degrees of purple, from highly to not significant (dark to light purple, respectively).

To extend the applicability of the machine learning models, we established the applicability domain based on the Tanimoto similarity network. Among the additional 1,563 antibiotics selected from PubChem, 569 (36.4%) with a Tanimoto similarity >0.2 compared to the 96 antibiotics used in this study were considered applicable (**Fig. 6b** and **Supplementary Data 7**). The best-performing models were used to predict the antibiotic resistance profiles stressed by these applicable antibiotics, resulting in resistance fold changes (log_2_ transformed) from −3.32 to 5.58 (**Fig. 6c** and **Supplementary Data 8**). To further evaluate the predictive accuracy of the models, we conducted experimental evolution using 16 external antibiotics (within the applicability domain and encompassing eight antibiotic classes, **Supplementary Table 1**). The predicted resistance levels to CIP and CAZ exhibited agreement with their actual values (*R*^2^_test_ > 0.75, **Supplementary Fig. 13**). These chemical structure-based models demonstrate promising potential for predicting antibiotic evolution under low-level antibiotic stresses.

In addition to ECFP modeling, we also explored mutation-based antibiotic resistance prediction. The AAC model was constructed based on the random forest (RF) algorithm, which indeed improved the prediction performance (*R*^2^_AAC_: 0.73 ± 0.05 vs. *R*^2^_ECFP_: 0.51 ± 0.15 for the training dataset, **Fig. 6a** and **Supplementary Fig. 14**). Despite proof-of-concept demonstrations, the applications of AAC-based modeling in predicting resistance phenotypes remain to be validated due to the complexity in intrinsic and extrinsic factors influencing mutation outcomes, e.g., genetic background and nutrient availability.

### Clinical and environmental prevalence of resistance-relevant mutations

Assessing the prevalence of key mutations in a broader range of clinical or environmental isolates is crucial for evaluating their risks in the ‘One Health’ paradigm. We focus on two groups of point mutations, comprising a total of 481 key SNPs, i.e., feature-importance SNPs contributing to the accuracy of the AAC-based RF models (**Fig. 6d**) and SNPs conferring high-level antibiotic resistance (**Supplementary Fig. 10**). These key SNPs and 1,263 *E. coli* genomes (clinical: 577, environmental: 686) retrieved from the NCBI Pathogen Detection database were subjected to comparative genomic analysis (**Supplementary Data 9 and 10**). Overall, 70 of the 481 key SNPs (14.6%) were identified in at least one of the external genomes (**Fig. 6e**), with the highest occurrence in 487 genomes (*yddB* Arg458His, **Fig. 6f**). Their relative prevalence in clinical and environmental isolates was summarized in **Supplementary Fig. 15**. Importantly, 62.9% of the key SNPs were identified in clinical isolates, and three were significantly more prevalent in clinical than environmental isolates, e.g., TetR family of transcription regulator gene *rutR* Ala15Thr and lipoprotein-related gene *bor* Ala66Thr. Furthermore, 74.3% of the key SNPs were identified in environmental isolates, and three were significantly more prevalent, e.g., oxidative stress response gene *btuE* Ala54Val. Notably, co-occurring SNPs (i.e., detected in both clinical and environmental isolates) accounted for 37.1% of the total key SNPs. These results suggested that mutations that significantly contribute to antibiotic resistance are widespread in both clinical and environmental isolates, which may be acquired under sub-MIC antibiotic exposure conditions.

## Discussion

Despite the growing attention that the development of AMR could be driven by the low-level selective pressure of antibiotics released into the environment, the relationship between AMR phenotypes and genotypes, and whether mutations conferring high-level resistance are prevalent remain key questions to be addressed. In this study, we constructed an unprecedented phenotype-genotype atlas of resistance evolution under 96 low-level antibiotic stress conditions. We used high-throughput experimental evolution and deep-resequencing approaches to determine, at the population level, how bacteria can evolve high-level resistance upon exposure to a wide range of antibiotics at low levels. Tn-Seq further facilitated the identification of PRGs and the elucidation of associated resistance mechanisms. ECFP- and AAC-based machine learning models have shown promise in areas such as predicting resistance phenotypes and identifying feature-importance antibiotic substructures and mutations. We leveraged an extensive collection of publicly available genomes to determine the prevalence of key mutations in a broader range of environmental and clinical settings. Some of the main findings extend beyond our understanding of resistance evolution under low-level antibiotic exposures, shedding insights into the origins and dissemination of antibiotic resistance in the environment.

From a phenotypic perspective, most stressors showed positive correlations in causing consistent antibiotic resistance changes in evolved populations. Cross-resistance was much more prevalent than collateral sensitivity^16^, reinforcing and complementing the previous findings of cross-resistance and collateral sensitivity relationships identified at clinical concentrations^36,37^. Some of the newly identified collateral sensitivity relationships (e.g., kanamycin or gentamicin vs. sparfloxacin, and aztreonam vs. ciprofloxacin; *P* < 0.05) offer valuable insights for developing alternative clinical antibiotic dosing strategies. The study is notable for pioneering the establishment of a link between stressor’s chemical structures and resistance evolution. For instance, quinolone stressors elicited the highest CIP resistance compared with other antibiotics, and their particular atomic groups, e.g., the C-6 fluorine atom and the cyclopropyl group, were found likely to contribute to high-level resistance evolution.

From a genotypic perspective, the results provide a comprehensive understanding of how critical mutations contribute to clinically relevant resistance and the evolutionary landscape. Frequently encountered ARGs such as *evgS* and *ermY* were among the most common targets of mutations under antibiotic stresses. Tn-Seq further enabled the construction of the GI network in response to antibiotics. We, therefore, identified a set of core mutant PRGs (*n* = 131) contributing to high-level resistance evolution through the phenotype-genotype atlas. In addition to the well-characterized ARGs, e.g., *pdeR* involved in biofilm formation^38^, *ycjV* involved in efflux pump^35,39^, *MJ1187* involved in drug inactivation^40^, *yafK* involved in target bypass^41,42^, and yjiR involved in transcriptional reprogramming^43^, others may confer high-level resistance via unreported mechanisms, e.g., *metH*, *ynfB*, *ulaG*, *ftsK*, *rtcR*, and *hflK*. By locating these understudied core mutant PRGs in the gene co-fitness interaction network and analyzing the functions of interacted genes, their resistance mechanisms were proposed. The resistance contribution mechanism of two core mutant PRGs, namely *rtcR* and *hflK*, through biofilm formation, was experimentally confirmed. Importantly, this demonstrated that the gene co-fitness network generated based on Tn-Seq data highlights valuable interactions that can be explored to uncover resistance mechanisms of potential ARGs. The SAFE analysis further suggests that genome-wide mutations affecting broad biological processes^44,45^, such as metabolism and cell motility, can significantly contribute to the evolution of antibiotic resistance^31,32^, extending beyond independent, target-gene-related mutations.

The ECFP- and AAC-based machine learning models exhibited promising levels of accuracy and scalability in elucidating the intricate relationships between resistance phenotype, genotype, and underlying drivers (i.e., antibiotics), leveraging the power of vast data that are unattainable through traditional low-throughput evolutionary experiments. Feature-importance antibiotic substructures identified may have important implications for the development of next-generation antibiotics and the targeted pollutant removal from contaminated environments^46^. The ECFP-based models exhibit broad applicability across a wide range of antibiotics, enabling relatively reliable prediction of antibiotic resistance in *E. coli* strains under low-level antibiotic exposures without the need for intensive, high-throughput experimental evolution. Moreover, the predicted resistance levels can be used as a reference, to some extent, for assessing the risks of environmentally acquired resistance.

Key SNPs selected at sub-MICs indeed exhibited both environmental and clinical relevance across a vast collection of *E. coli* isolates, and their environment-clinic co-occurrence rates were up to 37.1%. Some of these key SNPs were even enriched in clinical isolates, suggesting that the variants may already pose health risks. Consequently, antibiotic-contaminated aquatic and soil environments could serve as reservoirs of evolved resistant bacteria carrying clinically relevant, high-resistance-conferring mutations. A few studies have attempted to determine the genetic relationships between environmental and clinical isolates in public databases to infer transmission dynamics^47,48^. Our results made additional contributions to the identification of key mutations and ARGs of concern, as well as the quantification of their prevalence.

Disentangling the AMR risk is challenging and requires a holistic understanding of environment-clinic AMR dissemination, including but not limited to its evolution under low-level antibiotic exposures. Some considerations should be taken into account when extending our results to other studies on environmental resistance. First, environmental pathogens are remarkably diverse, likely resulting in species-specific response and resistance evolution patterns that differ from those of *E. coli*. Second, environmental changes are far more drastic and dynamic in nature, and resistance evolution is an ongoing process. Efforts to predict resistance profiles, either based on chemical structure or mutations, are inevitably constrained by temporal limitations and complex interplay of influencing factors. Third, uncovering the resistance-conferring mechanisms of core mutant PRGs is based on a hypothesis-verification approach. However, even if the hypotheses are experimentally validated, they may not be the primary mechanisms conferring resistance due to the pleiotropy of genes. We are actively investigating the extent to which these lab-scale findings from experimental evolution mirror AMR development in real-world settings, particularly antibiotic-contaminated environments. Nonetheless, these efforts provide valuable scientific support for combating antibiotic resistance within the One Health framework.

## Methods

### Bacterial strains, culture conditions, and antibiotics

*E. coli* strain QSHQ1 (GenBank accession number: CP128989) was a plasmid-free opportunistic pathogen that was originally isolated from the environment in Hangzhou, China (**Supplementary Fig. 16**). Cells were cultivated with LB medium (150 rpm, 37°C) in Abgene^TM^ 96-well 2.2 ml polypropylene deep-well plates (Thermo Scientific, #AB0661) with breathable sealing film (Corning, #1223S04). All antibiotics used were purchased from Yuanye-Bio, Macklin, and Solarbio (details in **Supplementary Data 1**). Antibiotic stock solutions (dissolved in 10% sterile DMSO) were filter-sterilized and stored at −20°C before use.

### Experimental evolution under 96 antibiotic stresses

Pre-adaptation: A single colony of *E. coli* strain QSHQ1 was picked from the LB-agar plate and grown in LB medium without antibiotics for 24 h. Cells were sub-cultured daily by 500-fold dilutions in fresh LB medium over a 10-day period, corresponding to approximately 9 generations per day.

Experimental evolution: 96 antibiotics were added to the LB-medium at a uniform concentration of 10 μg l^−1^. Four independent cultures (biological quadruplicate) were grown under each antibiotic stress and were sub-cultured daily by 500-fold dilutions for 20 days (1 ml medium, 96-well plates, 150 rpm, 37°C). The antibiotic concentration was selected to reflect the upper range of environmentally relevant concentrations detected in, e.g., wastewater treatment plants^49^. Four control lineages were cultured under the same condition without antibiotics. Cell growth was monitored by measuring OD_600_ using the Spark microplate reader (Tecan Inc.). After the 20-day experimental evolution, end-point cultures were frozen at −80°C in 30% glycerol for whole-genome sequencing. No growth was observed in the negative control wells throughout the experimental evolution.

### Automated high-throughput MIC assays of evolved populations

The automated high-throughput MIC assays were carried out on the iBioFoundry platform at ZJU-Hangzhou Global Scientific and Technological Innovation Center. The platform consisted of a MultiDrop Combi microplate dispenser (Thermo Scientific, #5840300), a Tecan Freedom Evo 200 Base laboratory automation workstation, a static incubator, and a CLARIOstar Plus microplate reader (BMG LABTECH Inc.), which were operated by two six-axis robotic arms (Thermo Scientific, TS-F7000).

Antibiotic resistance against azithromycin (AZI), ciprofloxacin (CIP), tetracycline (TET), sulfamethoxazole with trimethoprim (SMZ+TMP), tobramycin (TOB), ceftazidime (CAZ), meropenem (MER), and polymyxin B (PMB) were measured for evolved cultures at the end of the experimental evolution. The end-point glycerol-frozen cultures were revived in 1 ml LB medium for 16 h at 37°C with (evolved populations) or without (control) stressing antibiotics. The revived cultures were 2000-fold diluted in freshly prepared MIC plates to a final volume of 130 μl. The 2-fold serial dilutions were performed over 12 steps, starting from the highest concentrations of 400, 25, 100, 200, 100, 200, 25, and 400 mg l^−1^, respectively, for the eight antibiotics. After 20 h static incubation, the OD_600_ values were measured. The dose-response curves generated from biological quadruplicates were fitted to the Hill equation, and the antibiotic concentration corresponding to an OD_600_ of 0.2 was defined as IC_90_. In the event of curve overflow (exceeding 100% inhibition) or underfill (below 90% inhibition), the IC_90_ values were assigned as the maximum tested concentration or half of the minimum tested concentration, respectively.

### DNA extraction from evolved populations

In the antibiotic resistance assays, iBioFoundry identified the first well with detectable cell growth (OD_600_ > 0.2) in the presence of antibiotics, and pooled cells from up to 32 of these wells together (8 antibiotics × 4 replicates, ∼4 ml). Genomic DNA (gDNA) was extracted from pooled cultures using the DNeasy Blood & Tissue kit (Qiagen) in accordance with the manufacturer’s instructions. DNA quantity and quality were determined by the Colibri microvolume spectrometer (Titertek-Berthold Inc., Germany). High-quality DNA (A_260_/A_280_ = 1.8, > 6 μg) was used for whole-genome sequencing.

### Whole-genome sequencing and mutation analysis

Whole-genome sequencing was performed for the WT strain *E. coli* QSHQ1 as well as the 96 evolved populations, following the procedures described previously^10^. Briefly, the WT strain was sequenced using PacBio RS combined with Illumina sequencing (NovaSeq6000). The Illumina data were used to evaluate the complexity of the genomes and correct errors in PacBio long reads. The high-quality complete genome of the WT strain (5,084,285 bp) served as the reference for mutation detection (**Supplementary Fig. 17**). Pooled evolved populations were deep resequenced on the Illumina platform (150 bp×2, ∼1,500x coverage, Shanghai BIOZERON Co., Ltd). Mutations detected in the control lineages were excluded from further analysis. Variants with a frequency greater than 0.2% in the population were identified. The non-synonymous (d*N*) and synonymous (d*S*) substitutions were obtained under each stress condition, and the proportion of beneficial mutations was indicated by the *dN/dS* ratio, which was set to 1 for neutral evolution (without antibiotic stress).

### Transposon sequencing and gene fitness assays

A transposon mutant library for the WT strain was generated by electroporation using the EZ-Tn5 <KAN-2> Tnp transposome kit (Lucigen Inc.). The library consisted of approximately 40,000 insertion mutants, achieving a genome-wide saturation of 97.5%. The library was divided into 9 aliquots, 500x diluted in LB medium and grew for 24 h (4-8 generations) with no antibiotic or one of the eight antibiotics at sub-MIC levels. The gDNA was extracted, and a specific PCR forward primer (GATCCTCTAGAGTCGACCTGCAGG) was designed to amplify the regions flanking the Tn5 insertion (TA) sites. TA sites were sequenced on Illumina NovaSeq6000, and the sequencing data were analyzed using the TRANSIT package v3.2.7^50^. Essential genes were identified based on the Gumbel method by calling long consecutive sequences of TA sites without insertions. For non-essential genes with valid TA sites > 5 and mean insertion counts (*N*) > 5, roughly 3,500 in total, their gene fitness *W* was calculated as follows:

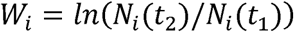

where *N*_i_(*t*_1_) and *N*_i_(*t*_2_) were the mean insertion counts of non-essential gene i detected in a particular library before and after antibiotic selection, respectively. The differences in the gene fitness were therefore calculated as Δ*W = W*_ABX_ – *W*_noABX_. Genes with absolute fitness difference values |Δ*W|* larger than 0.15 under at least one antibiotic stress condition were identified as potentially resistance-relevant genes (PRGs).

### Gene co-fitness interaction network and spatial analysis of functional enrichment (SAFE)

A co-fitness interaction network for PRGs was constructed as described previously^51^. In brief, a gene fitness matrix was built for a total of 2,830 PRGs, which was further used to calculate Pearson’s correlation coefficients (*r*) for all pairwise genes. A high-confidence co-fitness interaction network was established using Cytoscape, comprising 2,152 PRGs (nodes) and 22,811 interactions (edges) with |*r|* > 0.75 and corrected *P* < 0.00002 (threshold: 0.05, corrected by the number of genes). The plugin CytoHubba was used to calculate the degree and maximal clique centrality (MCC) of all nodes in the network. The plugin ClusterViz was employed to identify distinct modules based on network topology using the FAG-EC algorithm. PRGs were annotated using the Swiss-Prot, GO, CARD, VFDB, and BacMet databases. SAFE highlighted regions of the gene interaction network enriched in specific functional attributes.

### Knockout of selected PRGs and biofilm formation potential measurement

Two core mutant PRGs, *rtcR* and *hflK* were subjected to knockout experiments to verify their hypothesized involvement in biofilm formation. The single guide RNAs (sgRNAs) were designed using the online design tool (https://en.rc-crispr.com/). The pair of oligos for the two target genes were annealed and ligated to the CB-001 vector (Ubigene Biosciences Co., Ltd., Guangzhou, China). The CB-001-*rtcR* (or *hflK*) [gRNA] plasmids containing each target sgRNA sequence and donor were transformed into the WT strain by electroporation. Cells were spread onto LB agar and incubated overnight. Transformants were identified by colony PCR and Sanger sequencing. The sequences of sgRNAs and primers for CRISPR were provided in **Supplementary Tables 2** and **3**, respectively. Growth curves of the WT and knockout strains were obtained without antibiotics and with two low-level antibiotics (SMZ+TMP: 1 μg/ml, TOB: 5 μg/ml). The biofilm formation potentials of WT and knockout strains were determined based on the microtiter dish assay. In brief, strains were overnight grown in LB medium at 150 rpm and 37°C, and then diluted to OD_600_ = 0.05. The 96-well plates were statically incubated at 37°C for 24 h to allow biofilm formation, and the ultimate OD_600_ was determined. For staining, 200 μl of 0.1% crystal violet (CV) was added to each well and incubated for 15 min at 37°C. The wells were then washed three times with water and dried overnight. The stained biofilm was disrupted by adding 200 μl of 95% ethanol solution (95% EtOH, 4.95% distilled H_2_O, 0.05% Triton X-100), and OD_550_ was measured afterward. Biofilm formation potential was calculated as OD_550_/OD_600_.

### Machine learning methods

To elucidate the relationships between the chemical structures of antibiotics and the antibiotic resistance profiles they induced, quantitative structure-property relationship (QSPR) models were constructed based on a published study^52^. In brief, molecular fingerprints, including MACCS 166 and ECFPs and about 6,000 descriptors, were computed for the 96 stress-inducing antibiotics, which can be fine-tuned with a set of features. The antibiotic resistance profiles of evolved strains were used to train a number of ensemble regression models, including linear regression (OLS), partial least squares (PLS), k-nearest neighbors (KNN), and support vector machines (SVM). Automatic model generation (*n* = 10,000 iterations) by genetic algorithms was performed to find the best combination of features. The performance of a model was evaluated by using statistical parameters, i.e., in-sample coefficients of determination *R*^2^, RMSE, and out-of-sample random 10-fold cross-validation *R*^2^. The visualization strategy for tracing the atomic contribution was previously described^53,54^. To assess the applicability domain of the best-performing models, the structural data of 1,563 antibiotics, excluding the 96 used in experimental evolution, were downloaded from PubChem (https://pubchem.ncbi.nlm.nih.gov/#query=antibiotic). A pairwise Tanimoto similarity matrix was computed for the 1,659 antibiotics in total. An antibiotic was considered as inside domain if its Tanimoto similarity with at least one of the 96 antibiotics exceeded 0.2. The best-performing models were employed to predict antibiotic resistance development under stress conditions by the in-domain antibiotics. 16 out of the 569 in-domain antibiotics were randomly selected to evaluate model prediction performance, following the same procedures used in experimental evolution.

In addition to molecular fingerprint-based modeling, the relationships between 4,750 AACs identified in evolved strains and the resulting antibiotic resistance profiles were also explored using RF models. In brief, the Boole matrix was constructed based on the relative fold-change values and AACs for the 96 antibiotic stressors. The RF models employed 100 individual decision trees, each with a variable range of splits from 10 to 2,000 and a minimum split size of 5 per tree. Bootstrapping was employed to mitigate the inherent overfitting of models, where a 10-fold cross-validation (Shuffle Split) method was used for each machine-learning algorithm.

### Prevalence of point mutations based on comparative genomics

Comparative genomic analysis was performed using a well-established pipeline as described previously^31^. Briefly, all *E. coli* genomes with L50 < 20 were downloaded from the NCBI database (https://www.ncbi.nlm.nih.gov/pathogens/) on 3/24/2023, including ‘clinical’ (*n* = 3,376) and ‘environmental’ (*n* = 2,545) isolates. Genomes were searched for all key SNPs (*n* = 481) identified in this study. For each of the 5,921 genomes, Blast was used to identify the genomes hit for the QSHQ1 strain. If the BLAST e-value was 1*e*–10 or less, and the query genome had a coverage of at least 85%, the CDS sequences from hit clinical (*n* = 577) and environmental (*n* = 686) genomes were extracted, and multiple alignment was performed using Muscle. Mutated positions in all genomes in the database that aligned with the QSHQ1 SNPs were tabulated. Mutations that did not appear at least once in the entire pathogen database were removed from consideration; *P* values for over-representation in clinical (or environmental) isolates were calculated using Fisher’s exact test for all remaining mutations.

## Data availability

The raw sequence data of Reseq and Tn-Seq analyses are available in the NCBI Sequence Read Archive under the accession number PRJNA987173. All relevant data and models in this study are available from the corresponding authors upon reasonable request. Source data are provided with this paper.

## Acknowledgements

This work was supported by the ‘Major Program of National Natural Science Foundation of China (Grant No. 22193061 to Lizhong Zhu)’, the ‘Zhejiang Provincial Natural Science Foundation of China (Grant No. LR23B070002 to Huijie Lu)’, the ‘General Program of National Natural Science Foundation of China (Grant No. 22276165 to Huijie Lu)’, the ‘National Key Research and Development Program (Grant No. 2022YFC3704700 to Huijie Lu)’, and ZJU Foundation of Institute of Intelligent Biological and Chemical Manufacturing. The authors thank Dr. Haoxuan Liu from Zhejiang University Centre for Evolutionary & Organismal Biology and Dr. Lei Huang from ZJU-Hangzhou Global Scientific and Technological Innovation Center for valuable discussions.

## Author contributions

Conceptualization, H.W. and Huijie.L.; Software, H.W. and Hui.L.; Investigation, H.W. and Huijie.L.; Writing-original draft, H.W.; Writing-review & editing, H.W., Huijie.L., C.J. and L.Z.; Supervisions, Huijie.L. and L.Z. All authors approved the final paper.

## Materials & Correspondence

Correspondence and material requests should be addressed to Huijie Lu.

